# Engagement for Alcohol Escalates in the 5-Choice Serial Reaction Time Task After Intermittent Access

**DOI:** 10.1101/2023.11.30.569396

**Authors:** Phillip Starski, Addyson Siegle, Frederic Hopf

## Abstract

Uncontrollable binge drinking is becoming an increasingly prevalent issue in our society. This is a factor that plays a role in the development of alcohol use disorder (AUD). AUD impacts 15 million Americans annually, with approximately 88,000 dying from alcohol related deaths. There are several aspects of AUD that encourage a strong dependence on alcohol. Impulsivity, motivation, and attention are the primary behavioral facets we contribute to AUD. Many past studies have used the 5-Choice Serial Reaction Time Task (5-Choice) to analyze these types of behaviors using sugar as the reward. We have recently published a study where alcohol was used as a reward in the 5-Choice. 48 mice were trained to respond for alcohol in the 5-Choice, and the analyses for these animals were originally categorized by their alcohol preference and consumption. Upon looking at the data, we became more interested in a new way to classify these mice into groups. High engaged (HE) and low engaged (LE) mice were classified based on their number of correct responses in the last five late-stage sessions. During early-stage training, mice began to separate themselves into two groups based on their interaction with the task. The high-engaged (HE) mice were much more engaged with the task by having a high number of trials and correct responses, as well as a much lower percentage of omissions. The low engaged (LE) mice were not as engaged, this was apparent because of their lower number of trials and correct responses. They also had a much higher percentage of omissions in comparison to HE mice. LE mice presented no significant changes in late-stage training, while HE mice began responding and engaging more. These mice went through a period of intermittent access (IA), where they were allowed to drink alcohol in their cage for 3 weeks. After intermittent access, LE mice increased their responding which suggests an increase in motivation for alcohol as a reward. Engagement analysis presents two clearly different groups, one being motivated to work for alcohol and the other not wanting to work for this reward. These two distinct phenotypes in the 5-Choice could be used to model alcohol motivated behavior, which could help us further understand AUD.

## 1 INTRODUCTION

Alcohol Use Disorder (AUD) is a psychiatric condition that costs over $250 billion/year in the US, with majority cost stemming from those that binge drink ^1^. Excessive intake is strong feature of AUD than can contribute to the harmful effects of alcohol. This is especially in the case of enhancing the risk of drinking problems ^2-4^ and with the reduction of excessive intake decreases these risks and potential relapse ^5-7^. Additionally, those who do not have a dependency on alcohol but self-report greater impulsiveness have been seen to achieve higher blood alcohol levels in a free-access self-administration and experience greater euphoria from alcohol ^8^. The specific actions and cognitive processes an individual makes toward alcohol are potential points of investigation for potential treatment or biomarkers of future alcohol problems. One of these points is trait impulsivity, the innate general impulsiveness of an individual, which has been found to correlate with a greater risk of binge drinking ^9-11^. Importantly, impulsivity is a multi-faceted construct ^12-14^, with variants related to impulsive action (motor) and impulsive choice (cognitive) leading it to be seen as an important AUD risk factor. Another point of investigate is motivation for alcohol. Excessive intake leading to AUD triggers significant cognitive behavioral control changes ^15,16^. For example, the increased drive for intoxication and the developing tolerance with accompanied negative consequences (job loss, family troubles, or even jail time) display clear motivational changes in AUD individuals. ^17-19^. Lastly, AUD patients will develop an overt fixation on alcohol cues compared with ones for natural rewards suggesting an “attention bias” toward alcohol. ^20-22^. Ultimately, impulsivity, motivation, and attention make up the bulk of how an individual engages with alcohol and errors in this behavioral engagement processing, especially after chronic alcohol use, lead to harmful outcomes. Here, we seek to describe a novel way to investigate behavioral engagement for alcohol.

The five-choice serial reaction time task (5-Choice) can measure impulsive, attentional, motivational, and perseverative behavior across a single session ^23-25^. Further, the 5-Choice has strong translational value as it has been adapted for humans which has connected more impulsive individuals with greater alcohol intake ^26^. However, unlike human studies, rodent work allows for a deeper investigation into specific behavioral changes and further mechanistic insights. Studies examining the relation of alcohol and impulsivity have primarily determined 5-Choice responding for sugar in relation to alcohol exposure, until recently ^25-31^.

Here, we have re-analyzed data from our previous study that uniquely adopted the 5-Choice to have alcohol as the reward. We had found performance was significantly related to alcohol preference rather than consumption level (CITE). In this re-analysis, we identified a new classification called “engagement” that is based on correct responding of the last five sessions of late-stage training. This classification allows for a clearer phenotype of animals that actively work in the task for alcohol versus those that simply do not. Our previous analysis by preference (CITE) yields important information, however some animals who have high preference but do not engage in the 5-Choice (and vice versa) require further scrutiny. Using a median split of correct responding, we found strong correlation between preference and high engagement animals during early-stage training as well as clear differences in overall performance. Further, as in the previous analysis, we observed increased performance in low-engaged mice for alcohol after intermittent alcohol consumption, suggesting the vulnerability of individuals with initially lower drive for alcohol. Thus, we describe a new understanding of mouse 5-Choice behavior for alcohol that will be critical for interpretation of future brain mechanistic studies.

## 2 MATERIALS AND METHODS

### 2.1 Animals

Forty-eight male C57BL/6J mice from Jackson Laboratories Inc were individually housed, starting at 8 weeks old, in standard Plexiglass cages with *ad libitum* access to food and water until water restriction. Mice were maintained in a 12h:12h reverse light-dark cycle. Animal care and handling procedures were approved by the Indiana University Institutional Animal Care and Use Committee in accordance with NIH guidelines.

### 2.2 5-Choice serial reaction time task (5-Choice)

**(A) Water Restriction and Apparatus.** Water restriction was used throughout the experiment to promote performance for an alcohol reward ^32^. Water access was given for two hours after behavior sessions. Eight noise-attenuating Bussey-Sakisda Rodent Touch Screen Chambers (Lafayette Instruments Co., Lafayette, IN) were used for al behavior. These chambers have perforated stainless-steel floors with trapezoidal plastic walls enclosed with a touchscreen (12.1 in, resolution 800 x 600). Each screen had a plastic mask outlining five 4 x 4 cm holes spaced 1 cm apart and 1.5 cm from the floor. Infrared photo-detection beams were used to detect responses and movement within the chambers. Every chamber was also equipped with 3W lights, a speaker to deliver a 65 dB tone, ventilation fans, and an infrared-sensitive camera.
**(B) Pre-training.** (**Figure 1A**) Mice spent two days habituating to the operant chambers for 15 min where a 20ul reward of 10% alcohol was delivered after every tray exit to familiarize the mice to the reward location The following three days utilized a Fixed-Ratio 1 (FR1) schedule. Each of these sessions lasted 60 min with an unlimited number of trials available. Interestingly, Mice that completed the behavior earlier in the day performed less than those later in the day, so we added an additional FR1 session to reverse the order of the mice. Performance was unaffected.
**(C) Early-Stage Training.** (**Figure 1A**) Mice then began training in the 5-Choice. Mice are placed into the dark chamber are given a free 200ul reward (10% alcohol) that is illuminated in the reward tray in the rear of the arena. Each trial is initiated with a head entrance into the reward tray. Once initiated, the 5s intertrial interval (ITI) begins. Once the ITI ends, one of the 5 square holes illuminates for 10s (stimulus duration, SD). The animal has an additional 37s to give a response (limited hold, LH) before the trial is considered an omission. The animal can make several different responses. A touch on one of the five holes during the ITI period will result with a punishment (flash of light), recorded as a premature response, and will reset the ITI timer. If the stimulus is presented and the animal touches one of the unlit holes, there will be a punishment and it will be recorded as an incorrect response. After an incorrect or correct response is given, any additional responses in the five holes are recorded as a perseverative response. If a correct response is given, 20ul of 10% alcohol will dispense from a liquid pump in the illuminated reward tray (Figure 1B). After a correct, incorrect, or omission response, the reward tray will illuminate until the next tray exit, thereby initiating the next trial. Mice continued this training for 10 days.
**(D) Late-Stage Training.** (**Figure 1A**) As mice continued training, performance of both groups began to merge and classified this as late-stage training. Mice spent 14 days training in the 10s SD before being taken off water restriction to observe change in performance for 4 days. Next, the mice remained on ad libitum water access while being weight restricted to be tested for one week at 10s SD and 5s ITI for a sweet milk (Nesquik, Nestle) reward to compare performance between the 10% alcohol and a traditional 5-Choice reward.
**(B)** 5-Choice motivators: Water restriction was used because we wanted to heighten their response levels, and, importantly, in the oral modality that is used for alcohol; in other words, water restriction was more ethological as comparison for alcohol versus food restriction. For sweet milk sessions, as sweet milk is much more of a nutrient, food restriction was used.
**(C)** Post-Intermittent Access Testing. (**Figure 1A**). Four sessions for an alcohol reward and four for a sweet milk reward using a 10s SD and 5s ITI duration. Mice continued intermittent access alcohol intake across the testing weeks, resulting in approximately six weeks of total intermittent access, and were not weight restricted or continuously water restricted. Importantly, 5-Choice testing occurs after an intermittent access (IA) session. Briefly, a testing day begins with removal of the 24hr 20% alcohol bottle and no water access until, 5-9hrs later, we begin behavioral testing; this timing ensures mice were performing during acute withdrawal ^33,34^. Water bottles were given immediately after behavioral testing was completed for the day and remained until the next IA session. For analysis we excluded the first day to remove potential burst in behavior from reintroducing the mice to the task.

**Figure 1.**
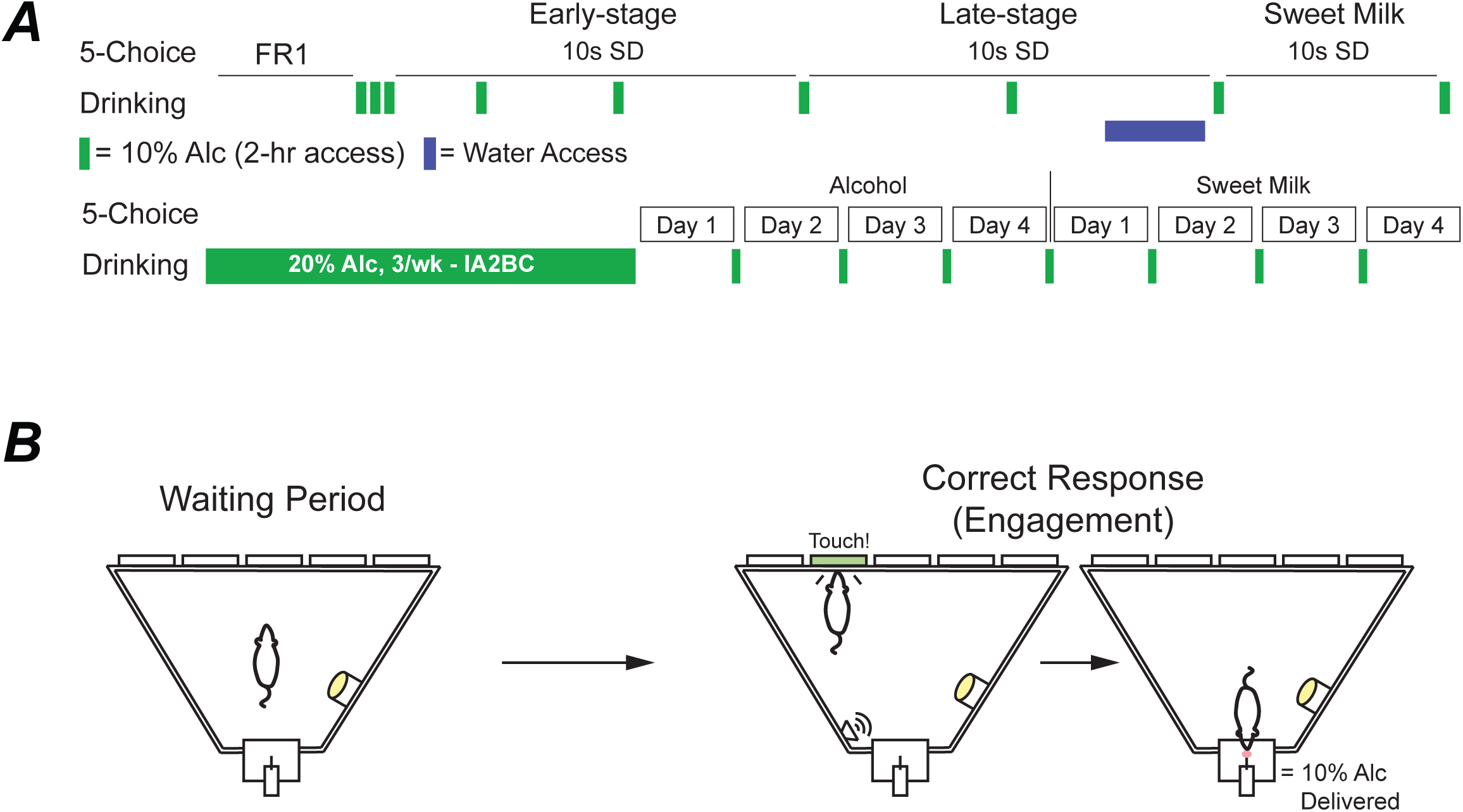
Experimental schematic. Timeline of the behavioral and drinking schedule (A). Example cartoon of the 5-Choice waiting period (5s) and a correct response that is used to determine engagement in the task (B).

### 2.3 Drinking in the Dark and Intermittent Access

**(A)** Throughout 5-Choice training, mice were given weekend drinking in the dark (DID) session of 10% alcohol for 2 h to promote response to the reward (**Figure 1A**). Custom-built, low drip sipper tubes were used to reduce dripping from overactive mice that may climb on the cage. These tubes consisted of a Falcon 15 mL conical tube (Fisher Scientific, Hampton, NH) with the bottom cone cut and filed down. A sipper (Ancare Corp, Bellmore, NY) was then placed inside the tube and shrink-wrapped using a heat gun. A rubber stopper (size:0#, StonyLab, Nesconset, NY) was used to plug the opposite end. Bottles were weighed before and after sessions and consumption was calculated using the weekly weight of the mouse. Preference was calculated as the total amount of alcohol consumed divided by total liquid consumed.
**(B)** Similar to Lei et al. 2019 ^35^, mice were given 24 h access to 20% alcohol every Sunday, Tuesday, and Thursday starting at 7:00 am (**Figure 1A**). Consumption was calculated using the weekly weight of the mouse. During non-alcohol days, two-bottles filled with water were present to maintain familiarity with the bottles.

### 2.4 Statistical Analysis

48 mice were categorized as High/Low preference, consumption, and engagement after the Late-Stage sessions and once the DID behavior was completed. Specifically, alcohol consumption and preference were calculated for each animal based on the overall average of the nine DID sessions. For classifying preference groups, mice were ordered from greatest preference to least preference and a median split was used to divide the 48 mice into two equal-sized groups. For classifying consumption groups, the same mice were instead ordered from greatest consumption to least consumption and a median split was used to divide mice into two equal-sized groups. For engagement, the median split was based on the average number of correct responses during the baseline sessions (last five late-stage training sessions). 5-Choice and alcohol intake studies were analyzed by two-way repeated measures analysis of variance (ANOVA) followed by Bonferroni’s multiple comparisons test where appropriate. Sphericity was not assumed, and the Geisser-Greenhouse correction was used. For missing data points (spill during drinking, or animal non-responding), a mixed-model analysis of variance was used. All group analyses of pre-IA versus post-IA and performance changes were tested for normal distribution (Shapiro-Wilk test), and then an appropriate test (parametric or non-parametric) was used to measure differences (t-test, Mann-Whitney, Paired t-test, Wilcoxon). All statistical analyses were calculated using Prism 9.0 software (Graphpad Software Inc., San Diego, CA), with significance set at *p*<0.05.

## 3 RESULTS

### 3.1 High- and low-engaged mice display similar alcohol preference and consumption but high engagement have greater performance during Early- and Late-stage training

Similar to our previous analysis (CITE), we used alcohol drinking and preference calculated from the first nine 2 hour-DID two-bottle-choice sessions (**Figure 1A**). To classify mice as either high-engaged (HE, n=24) or low-engaged (LE, n=24) we used a median split of the average correct responses during baseline sessions where HE averaged greater and LE averaged less than the median (16 correct responses). HE and LE displayed similar alcohol preference (**Figure 2A**, HEvLE: F1_1,46_=1.583, *p*=0.2147; time: F_5.519,251.8_=2.050, *p*=0.0654) and consumption (**Figure 2B**, HEvLE: F_1,46_=0.5712, *p*=0.4536; time: F_6.270,286.8_=17.73, *p*<0.001). **Figure 2C** and **D** detail the distribution of high/low preference and consumption within HE and LE groups.

**Figure 2.**
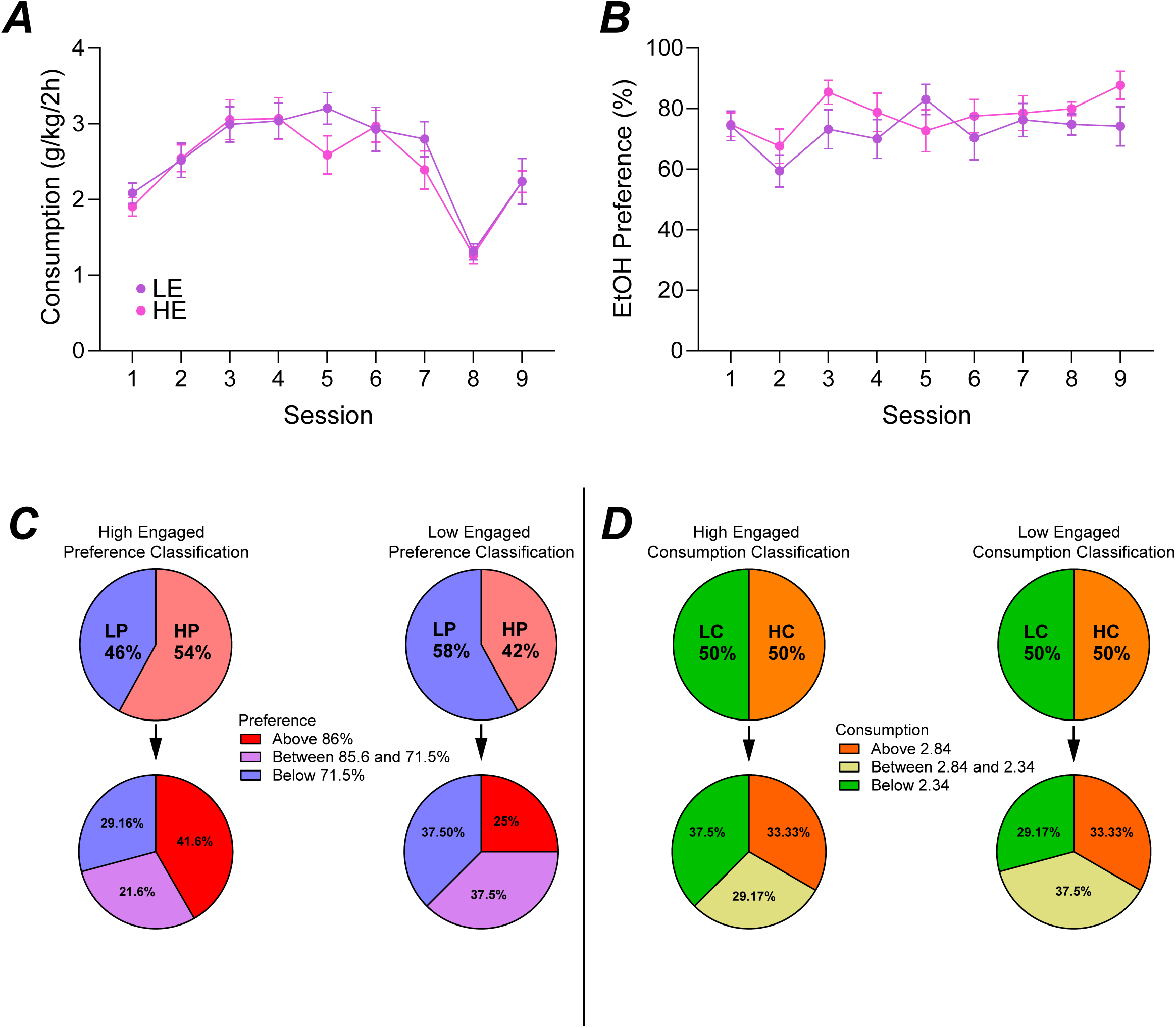
Animal classification based on engagement. High-engaged (HE) and low-engaged (LE) had similar alcohol consumption (A) and preference (B) during homecage drinking sessions. Previously, these animals were classified as either high/low-preference or consumption for analysis. The amount of HE and LE mice previously considered high-preference or low-preference with further separation based on range of preference (C). The amount of HE and LE mice previously considered high-consumption or low-consumption with further separation based on range of consumption (D).

Mice were trained in the 5-Choice for a 10% alcohol reward and the first ten days (Early-stage) were analyzed to observe the initial motivation to learn the task. HE mice showed significantly greater number of trials (**Figure S1A**, HEvLE: F_1,46_=23.04, *p<*0.0001; time: F_2.119,97.45_=9.588, *p*<0.0001, accuracy (**Figure 3A_1_,** HEvLE: F_1,46_=6.758, *p*<0.0125; time: F_4.828,222.1_=6.602, *p*<0.0001; interaction: F_9,414_=2.193, *p*=0.0217, with post-hoc significance on sessions 10), correct responses (**Figure 3B_1_,** HEvLE: F_1,46_=17.78, *p*<0.0001; time: F_1.800,82.81_=8.862, *p*=0.0005; interaction: F_9,414_=6.701, *p*<0.0001, with post-hoc significance on sessions 2, 5, 8 through 10), premature responses (**Figure S1B,** HEvLE: F_1,46_=12.94, *p*=0.0008; time: F_1.430,65.78_=4.189, *p*=0.0311; interaction: F_9,414_=3.124, *p*=0.0012, with post-hoc significance on sessions 1, 4-8), percentage of premature responses (**Figure S1C,** HEvLE: F_1,46_=21.71, *p*<0.0001; time: F_3.205,147.4_=2.980, *p*=0.0304; interaction: F_9,414_=4.027, *p*<0.0001),with post-hoc significance on sessions 1, 2, 4, and 6), and lower percentage of omissions (**Figure 3A_3_,** HEvLE: F_1,46_=31.95, *p*<0.0001; time: F_3.767,173.3_=1.964, *p*=0.1062; interaction: F_9,414_=8.892, *p*<0.0001, with post-hoc significance on sessions 4-10) compared to LE mice. As the classification of mice are based on their baseline sessions (last five sessions during Late-stage), this data may suggest a subset of mice have an innate drive to perform for an alcohol reward.

**Figure 3.**
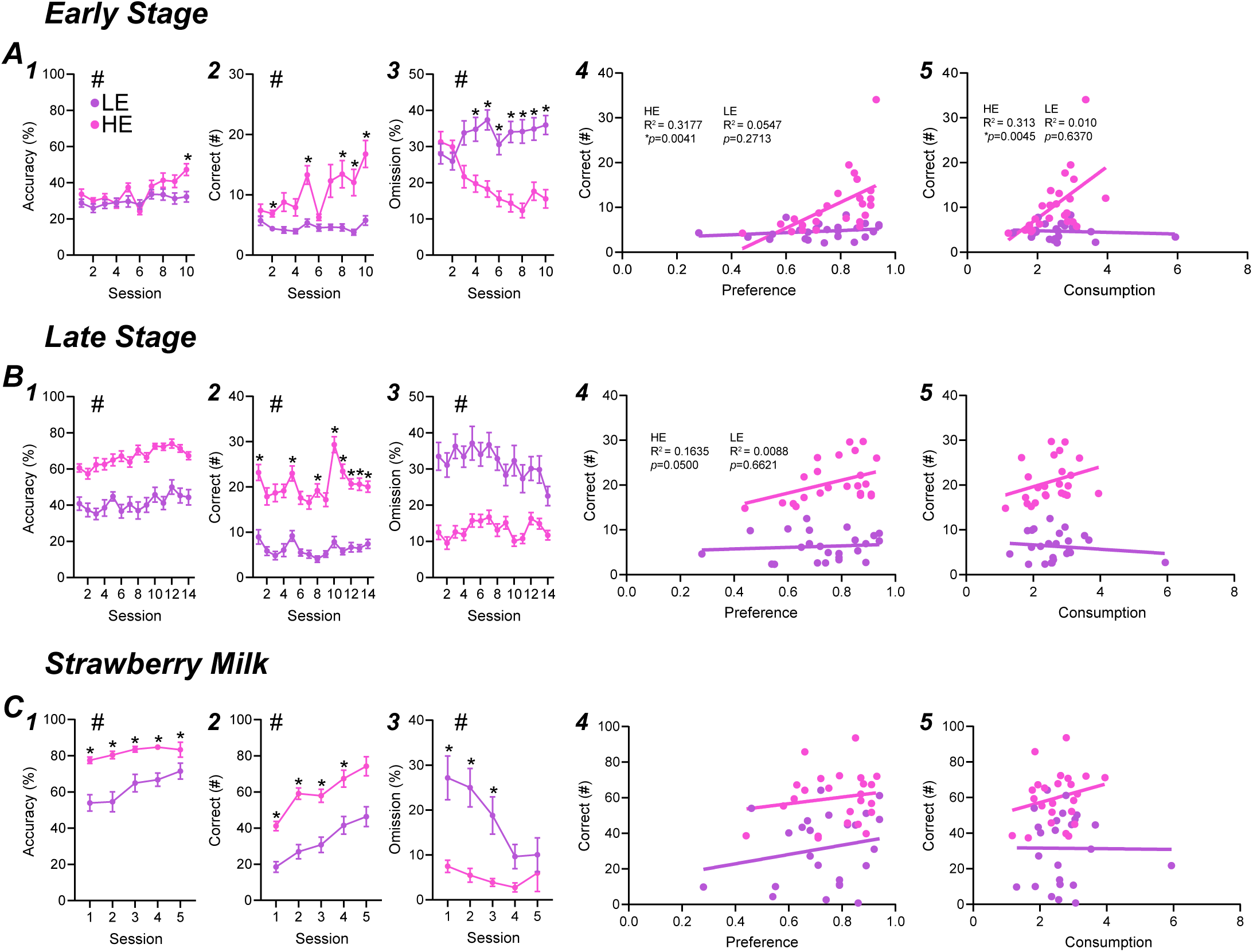
High-engaged mice perform greater during all phases of training. Early stage performance was much greater in HE mice with increased accuracy, correct responding, and decreased omissions. Additionally, HE mice correct responding correlated with homecage alcohol preference and consumption (A). HE continued to have greater behavioral performance during late stage training with a strong correlative trend of HE correct responding to homecage alcohol preference (B). When the reward was switched to strawberry milk, both groups greatly increased performance with HE mice performing greater. LE mice, if tested longer, are hypothesized to match HE performance as described by the significant decrease in omission percentage (C).

Training continued for an additional 14 sessions to see if mice could be adequately trained in the 5-Choice with an alcohol reward (Late-Stage). HE mice showed significantly greater number of trials (**Figure S1D,** HEvLE: F_1,46_=98.75, *p*<0.0001; time: F_7.843,360.8_=19.64, *p*<0.0001; interaction: F_13,598_=2.623, *p*=0.0015), with post-hoc significance on sessions 1-14), accuracy (**Figure 3B_1_,** HEvLE: F_1,46_=61.85, *p*<0.0001; time: F_7.364,338.7_=5.392, *p*<0.0001), correct responses (**Figure 3B_2_,** HEvLE: F_1,46_=136.9, *p*<0.0001; time: F_7.531,346.4_=9.463, *p*<0.0001; interaction: F_13,598_=3.271, *p*<0.0001), with post-hoc significance on sessions 1-14), premature responses (**Figure S1E,** HEvLE: F_1,46_=31.68, *p*<0.0001; time: F_5.172,237.9_=11.62, *p*<0.0001; interaction: F_13,598_=5.983, *p*<0.0001), with post-hoc significance on sessions 5, 9, and 10), percentage of premature responses (**Figure S1F,** F_1,46_=14.77, *p*=0.0004; time: F_9.029,415.4_=5.713, *p*<0.0001; interaction: F_13,598_=3.417, *p*<0.0001),with post-hoc significance on sessions 1, 3, 5, 8-11), and lower percentage of omissions (**Figure 3B_3_,** HEvLE: F_1,46_=51.55, *p*<0.0001; time: F_7.132,328.1_=3.098, *p*=0.0033) compared to LE mice. HE mice consistently out-performed LE mice during late-stage training and emphasize the phenotypic difference in motivation between mice responding for alcohol.

Baseline sessions consisted of the last five sessions of Late-stage training and were the basis for the median split. HE mice showed significantly greater number of trials (**Figure S1D,** HEvLE: F_1,46_=116.4, *p*<0.0001; time: F_3.229,148.5_=12.88, *p*<0.0001), accuracy (**Figure 3B_1_,** HEvLE: F_1,46_=57.24, *p*<0.0001; time: F_3.466,459.4_=2.212, *p*=0.0795), correct responses (**Figure 3B_2_,** HEvLE: F_1,46_=195.3, *p*<0.0001; time: F_2.966,136.4_=8.954, *p*<0.0001; interaction: F_4,184_=6.329, *p*<0.0001, with post-hoc significance on each session), premature responses (**Figure S1E,** HEvLE: F_1,46_=29.39, *p*<0.0001; time: F_1.648,75.81_=17.94, *p*<0.0001; interaction: F_2.966,136.4_=8.954, *p*<0.0001, with post-hoc significance on sessions 1 and 2), percentage of premature responses (**Figure S1F,** HEvLE: F_1,46_=5.712, *p*=0.021; time: F_3.247,149.4_=8.377, *p*<0.0001; interaction: F_4,184_=8.303, *p*<0.0001, with post-hoc significance on session 1), and lower percentage of omissions (**Figure 3B_3_,** HEvLE: F_1,46_=27.70, *p*<0.0001; time: F_3.495,160.8_=3.224, *p*=0.0185) compared to LE mice. Even after 24 sessions, HE mice remain greatly more motivated to respond for alcohol compared to LE mice.

### 3.2 High alcohol preference and consumption is predictive of early-stage performance

To find potential relationships between engagement behavior and alcohol drinking, we compared the average number of correct responses during Early-, Late-, and baseline sessions to alcohol preference and consumption.

For alcohol preference comparison, there was an overall (n=48) correlation during early-stage (**Figure S2A,** r^2^=0.1683, *p*=0.0038), but not late-stage (**Figure 2B**, r^2^=0.2625, *p*=0.0714) or baseline sessions (**Figure 3C**, r^2^=0.2115, *p*=0.1490). When comparing HE and LE to alcohol preference separately, we observe a strong correlation between early-stage and HE (**Figure 3A_4_,** r^2^=0.3177, *p*=0.0041) mice, but not LE (**Figure 3A_4_,** r^2^=0.0547, *p*=0.2713). During Late-stage, HE held a strong trend in relation to preference (**Figure 3B_4_,** r^2^=0.1635, p=0.0500) opposed to LE (**Figure 3B_4_,** r^2^=0.0088, *p*=0.6621). HE (**Figure S2D,** r^2^=0.0847, *p*=0.1676) and LE (**Figure S2D,** r^2^>0.0001, *p*=0.9787) did not show any correlation to preference during baseline sessions. For alcohol consumption comparison, there was no overall (n=48) correlation between Early-stage (**Figure S2E,** r^2^=0.0351, *p*=0.2021), Late-stage (**Figure S2F,** r^2^=0.6706, *p*=0.0039), or baseline sessions (**Figure S2G,** r^2^=0.0079, *p*=0.5467). When comparing HE and LE to alcohol consumption separately, we observe a strong correlation between early-stage and HE (**Figure 3A_5_,** r^2^=0.3127, *p*=0.0045) mice, but not LE (**Figure 3A_5_,** r^2^=0.0103, *p*=0.6370). During late-stage, HE (**Figure 3B_5_,** r^2^=0.0939, *p*=0.1453) and LE (**Figure 3B_5_,** r^2^=0.0202, *p*=0.5073) did not show any correlation during late-stage training. HE (**Figure S2H,** r^2^=0.03704, *p*=0.3676) and LE (**Figure S2H,** r^2^=0.0105, *p*=0.6336) did not show any correlation to preference during baseline sessions. These data suggests that alcohol preference and consumption predicts early motivation in the 5-Choice for alcohol, where preference may potentially predict greater performance in later training.

### 3.3 Sweet milk performance escalated significantly in both groups

Next, to determine behavioral differences between an alcohol reward and a traditional sugar-based reward (sweet milk), mice were no longer water restricted, but weight restricted (Methods and Discussion further explain this reasoning). The sweet milk 5-Choice behavior lasted five sessions. HE mice displayed overall greater number of Trials (**Figure S1G,** HEvLE: F_1,45_=26.59, *p*<0.0001; time: F_2.789,125.4_=41.75, *p*<0.0001), correct (**Figure 3C_2_,** HEvLE: F_1,45_=31.78, *p*<0.0001; time: F_2.668,119.4_=53.28, *p*<0.0001), accuracy (**Figure 3C_1_,** HEvLE: F_1,45_=21.58, *p*<0.0001; time: F_3.147,141.6_=9.966, *p*<0.0001; interaction: F_4,180_=2.698, *p*=0.0323, with post-hoc significance on sessions 1-4), percent omission (**Figure 3C_3_,** HEvLE: F_1,45_=14.33, *p*=0.0005; time: F_2.696,121.3_=8.347, *p*<0.0001; interaction: F_4,180_=4.932, *p*=0.0009, with post-hoc significance on sessions 1-3), premature (**Figure S1H,** HEvLE: F_1,45_=13.37, *p*=0.0007; time: F_3.367,151.5_=4.513, *p*=0.0032), but no difference in percent premature (**Figure S1I,** HEvLE: F_1,45_=2.335, *p*=0.1335; time: F_3.321,148.6_=1.213, *p*=0.3078). It is clear that HE mice perform better in the 5-Choice, however LE responding escalated each subsequent session. We hypothesize that since HE mice were already more active in the 5-Choice when responding for alcohol, then they have had additional practice toward maximizing their responding. In contrast, LE mice spent much less time responding in the 5-Choice for alcohol and may need additional time to develop their skill in the task. We would expect that LE mice would converge with HE mice if more sweet milk sessions were completed.

### 3.4 High-engaged mice display greater motivation for the reward, especially for alcohol

Motivation for the reward is a critical aspect of behavioral engagement. In the 5-Choice, the time between a correct response and retrieval of the reward describes the animal’s motivation for the reward. Here, using alcohol, we see a clear difference between HE and LE mice during Early-Stage (**Figure 4A**, HEvLE: F_1,46_=18.98, *p*<0.0001; time: F_3.219,148.1_=1.882, *p*=0.1308), but collapsed as sessions progressed to the Late-Stage (**Figure 4B**, HEvLE: F_1,46_=1.594, *p*=0.2131; time: F_6.244,286.7_=2.206, *p*=0.0402). However, HE mice showed greater motivation during sweet milk sessions (**Figure 4C**, HEvLE: F_1,45_=5.290, *p*=0.0261; time: F_2.890,127.9_=1.867, *p*=0.1405; interaction: F_4,177_=4.314, *p*=0.0024, with post-hoc significance on sessions 1, 2 and 5). Further, Baseline (**Figure 4A**, sessions 11-15) sessions were averaged across the five sessions. LE mice displayed greater latency than HE mice (*U =* 183, *p*=0.0483) for alcohol as well as for sweet milk (*U =* 146, *p*=0.0086). Additionally, LE were greatly more motivated for sweet milk than for alcohol (*U =* 110, *p*=0.0003) and similarly found in HE mice (*U =* 82, *p*=<0.0001).

**Figure 4.**
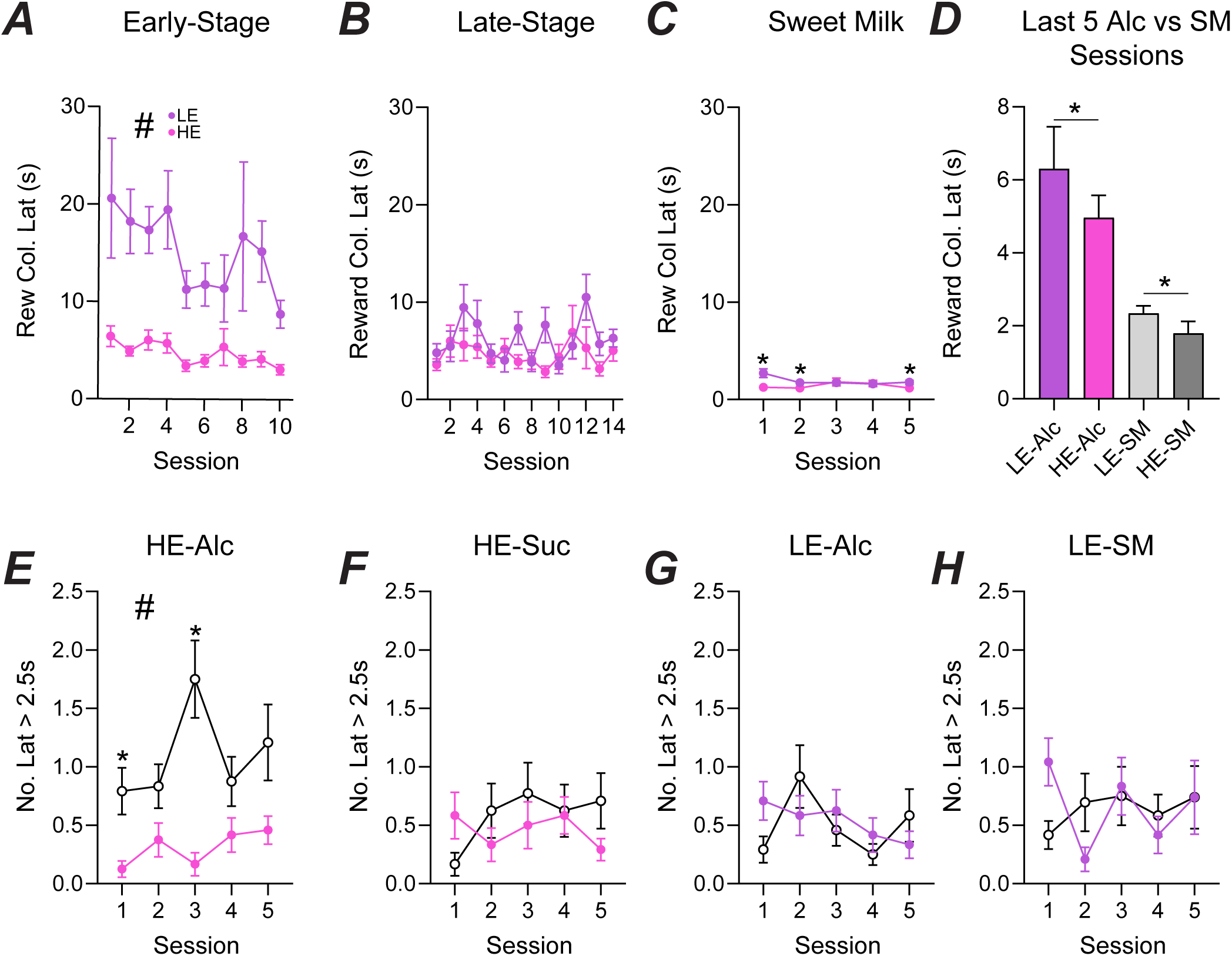
HE mice show high motivation for an alcohol reward primarily during the first half of the session. HE mice displayed significantly lower reward latency during Early (A) training, but both HE and LE were similar during late stage (B) and sweet milk training (C). HE mice had significantly lower reward latency during the final five sessions of late stage and sweet milk training (D). The number of reward latencies greater than 2.5s occurred more frequently during the last half of the session for HE mice for an alcohol reward (E) unlike HE for sweet milk (F), LE for alcohol (G), and LE for sweet milk (H).

As alcohol is an intoxicant, we are concerned that the mice will face motor deficits as they continue to consume alcohol throughout the session. Thus, we decided to separate their session into two halves. The trials for the HE mice during alcohol sessions showed that most of their reward latencies above 2.5 seconds occurred in the second half (**Figure 4E**, HEvLE: F_1,46_=25.36, *p*<0.0001; time: F_3.274,150.6_=2.181, *p*=0.0870; interaction: F_4,184_=3.045, *p*=0.0184, with post-hoc significance on sessions 1 and 3). No differences were found for HE during sweet milk (**Figure 4F**, HEvLE: F_1,46_=0.8197, *p*=0.3700; time: F_3.478,158.3_=0.6155, *p*=0.6296), nor for LE during alcohol (**Figure 4G**, HEvLE: F_1,46_=0.0662, *p*=0.7980; time: F_3.470,159.6_=1.847, *p*=0.1318) and LE during sweet milk sessions (**Figure 4H**, HEvLE: F_1,46_=0.0010, *p*=0.9748; time: F_2.819,127.6_=1.273, *p*=0.2865). As the increased in longer latencies occurred only in HE mice during alcohol, we can hypothesize that alcohol does play a role in behavior during the 5-Choice. However, the importance of this role will need to be explored further.

### 3.5 Low-engaged mice significantly reduce reward collection latency after intermittent access

During phase 2, all mice underwent intermittent alcohol access where they were exposed to 20% alcohol for 24hr three times a week. Mice were retested in the 5-Choice for four sessions with an alcohol reward followed by four sessions with a sweet milk reward. Importantly, as the first session causes a burst of behavior as we described in the previous paper (CITE), we only used the last three days for analysis. Further, to compare change in behavior from pre-intermittent access (IA) to post-IA, we only used the last three days of the Late-Stage and sweet milk sessions.

First, we made reward collection latency comparisons by engagement behavior (summarized in **Table 1**). Having only three pre-IA sessions, LE mice still displayed greater reward collection latency compared to HE mice for alcohol (Mann-Whitney, *U*=172, *p*=0.0266) and for sweet milk (Mann-Whitney, *U*=167.5, *p*=0.0202). Interestingly, after IA, LE and HE mice displayed similar latencies (Mann-Whitney, *U*=240, *p*=0.3305) for alcohol, but sweet milk remained different (student’s t-test, t[45]=3.914, *p*=0.0003). More specifically, LE mice reward latency significantly greater pre-IA than post-IA (Wilcoxon test: *p*=0.0079) whereas HE mice showed no significant change (Wilcoxon test: *p*=0.2897). Additionally, LE (paired t-test, t[22]=2.589, *p*=0.0168) decreased latency for sweet milk after IA and HE held a strong trend decrease (paired t-test, t[23]=1.938, *p*=0.0650). The overall change in performance from pre-IA to post-IA was significantly greater in LE mice compared to HE mice (student’s t-test, t[46]=2.315, *p*=0.0252). However, there was no difference in pre/post-IA sweet milk performance between HE and LE mice (student’s t-test, t[45]=0.0753, *p*=0.9402).

When analyzing by alcohol preference (High-preference, HP, and Low-preference, LP), we observed that LP mice displayed similar reward collection latency compared to HP mice for alcohol (Mann-Whitney, *U*=283, *p*=0.9268) and for sweet milk (Mann-Whitney, *U*=239.5, *p*=0.4441). Interestingly, after IA, HP mice displayed faster reward latencies (Mann-Whitney, *U*=164, *p*=0.0100) for alcohol, but not for sweet milk (student’s t-test, t[45]=0.1881, *p*=0.8516). More specifically, LP mice reward latency significantly greater pre-IA than post-IA (Wilcoxon test: *p*=0.0126) but HP mice did not change post-IA (Wilcoxon test: *p*=0.2076). Additionally, LP (Wilcoxon test: *p*<0.0001) significantly decreased latency for sweet milk post-IA while HP (paired t-test, t[22]=1.918, *p*=0.0681) held a strong trend toward decreased latency. The overall change in performance from pre-IA to post-IA was not different between LP and HP mice (student’s t-test, t[46]=0.3910, *p*=0.6976). However, there was no difference in pre/post-IA sweet milk performance between HP and LP mice (student’s t-test, t[45]=0.3345, *p*=0.7396).

When analyzing by alcohol consumption (High-consumption, HC, and Low-consumption, LC), we observed that LC and HC mice had similar reward collection latency for alcohol (Mann-Whitney, *U*=285, *p*=0.9552) and for sweet milk (Mann-Whitney, *U*=260.5, *p*=0.7480). Interestingly, after IA, HC mice displayed slower reward latencies (Mann-Whitney, *U*=177, *p*=0.0352) for alcohol compared to LC, but both remained similar for sweet milk (Mann-Whitney, *U*=265, *p*=0.8248). More specifically, LC mice reward latency significantly greater pre-IA than post-IA (Wilcoxon test: *p*=0.0079) but HC mice did not change post-IA (Wilcoxon test: *p*=0.2643). Additionally, LC (Wilcoxon test: *p*<0.0001) and HC (Wilcoxon test: *p*=0.0179) both decreased latency for sweet milk post-IA. The overall change in performance from pre-IA to post-IA was not different between LC and HC mice (student’s t-test, t[46]=0.2021, *p*=0.8407). There was also no difference in pre/post-IA sweet milk performance between HC and LC mice (student’s t-test, t[45]=0.1014, *p*=0.9197).

Reward collection latency is critical for identifying motivation for the reward. Here, engagement analysis shows that LE mice are significantly less motivated for the reward than HE mice prior to IA. Interestingly, LE mice become greatly motivated for alcohol to similar levels of HE mice after IA. By separating animals by performance in the 5-Choice, not preference or consumption, we observe a greater behavioral change caused by IA. As LE mice became greatly motivated for alcohol after excessive alcohol exposure, this may suggest that low-engaged individuals are still susceptible to alcohol issues.

### 3.6 Stimulus response latency of low-engaged mice changes significantly more than high-engaged mice after intermittent access

Next, we wanted to further characterize the motivation of the mice during the task. Thus, we compared correct and incorrect latencies. This is the time from when the intertrial interval waiting period ends and the time it takes them to give a response. A faster latency indicates greater motivation to perform the task. We found that HE mice were significantly faster than LE mice during pre-IA alcohol (student’s t-test, t[46]=0.4329, *p<*0.0001) and sweet milk session (student’s t-test, t[45]=2.917, *p*=0.0055). However, after IA, LE and HE mice displayed similar correct latency for alcohol (student’s t-test, t[46]=4.309, *p*=0.6685), but still showed a difference in sweet milk (student’s t-test, t[45]=2.715, *p*=0.0094). LE mice were significantly faster to give a correct answer for alcohol post-IA compared to pre-IA (paired t-test, t[23]=3.220, *p*=0.0038), as did HE mice (paired t-test, t[23]= 2.837, *p*=0.0093). However, LE had a greater change in latency than HE mice (student’s t-test, t[46]=4.081, *p*=0.0002). For sweet milk, LE (paired t-test, t[23]=2.144, *p*=0.0429) and HE (paired t-test, t[23]=2.871, *p*=0.0089) were significantly faster to give a correct response after IA compared to pre-IA, but there was no significant difference in the change in performance between them (student’s t-test, t[45]=1.304, *p*=0.1988).

When comparing by alcohol preference, we found that HP and LP mice had similar correct latencies during pre-IA alcohol (student’s t-test, t[46]=1.547, *p=*0.1287) and sweet milk session (student’s t-test, t[45]=1.831, *p*=0.0737). After IA, LP and HP mice also displayed similar correct latency for alcohol (student’s t-test, t[46]=0.9402, *p*=0.3520) and no difference when responding for sweet milk (student’s t-test, t[45]=1.232, *p*=0.2243). LP mice were significantly faster to give a correct answer for alcohol post-IA compared to pre-IA (paired t-test, t[23]=2.301, *p*=0.0308), but HP mice did not (paired t-test, t[23]= 0.2828, *p*=0.7798). However, LP had a strong trend toward a greater change in latency than HP mice (student’s t-test, t[46]=1.984, *p*=0.0532). For sweet milk, LP (paired t-test, t[23]=1.319, *p*=0.2003) and HP (paired t-test, t[23]=1.690, *p*=0.8674) did not show any difference in latency after IA compared to pre-IA, and there was no significant difference in the change in performance between them (student’s t-test, t[45]=1.116, *p*=0.2702).

Consumption analysis found that HC and LC mice had similar correct latencies during pre-IA alcohol (student’s t-test, t[46]=0.8319, *p=*0.4098) but HC were significantly faster during sweet milk sessions (student’s t-test, t[45]=2.377, *p*=0.0218). After IA, LC and HC mice also displayed similar correct latency for alcohol (student’s t-test, t[46]=0.1564, *p*=0.8764) and maintained their difference when responding for sweet milk (student’s t-test, t[45]=2.102, *p*=0.0411). LC mice responded similarly for alcohol post-IA compared to pre-IA (paired t-test, t[23]=1.555, *p*=0.1336), as did HC mice (paired t-test, t[23]= 0.5654, *p*=0.5773). However, LC had similar changes in latency than HC mice (student’s t-test, t[46]=0.9042, *p*=0.3706). For sweet milk, LC (paired t-test, t[23]=2.144, *p*=0.0429) decreased in correct latency after IA in contrast to HC (paired t-test, t[23]=0.8096, *p*=0.4268). There was no significant difference in the change in performance between HC and LC mice (student’s t-test, t[45]=1.288, *p*=0.2042).

Further, we wanted to look at the average time when giving an incorrect response. Mice with low incorrect response rates could potentially support any cognitive errors in impulse control or attention. Pre-IA, HE and LE mice had similar incorrect latencies (student’s t-test, t[46]=1.002, *p*=0.3214) as well as post-IA (student’s t-test, t[46]=0.7406, *p*=0.4627) when responding for an alcohol reward. LE mice did not significantly change performance post-IA (paired t-test, t[23]=0.3807, *p*=0.7069), however HE mice significantly increased post-IA (paired t-test, t[23]=2.414, *p*=0.0241). The overall change in performance between HE and LC mice were not significantly different (student’s t-test, t[46]=1.286, *p*=0.2050). For sweet milk, HE mice had faster incorrect latencies pre-IA (t[45]=3.231, *p*=0.0023) and post-IA (t[45]=2.408, *p*=0.0202) compared with LE mice. Both HE (paired t-test, t[23]=0.2053, *p*=0.8391) and LE (paired t-test, t[22]=1.613, *p*=0.1209) did not significantly change performance post-IA. The overall change in performance between HE and LE mice were not significantly different (student’s t-test, t[45]=1.292, *p*=0.2029).

Preference analysis yielded no differences pre-IA, as HP and LP mice had similar incorrect latencies (student’s t-test, t[46]=0.8503, *p*=0.3996) for alcohol and sweet milk (student’s t-test, t[46]=1.104, *p*=0.2755). After IA, there were no differences in incorrect latency for alcohol (student’s t-test, t[46]=0.1221, *p*=0.9034) or sweet milk (student’s t-test, t[45]=0.0736, *p*=0.9416) between HP and LP mice. HP incorrect latency did not change after IA when responding for alcohol (paired t-test, t[23]=1.736, *p*=0.0959) or sweet milk (paired t-test, t[22]=0.0390, *p*=0.9692). LP mice also saw no changes in incorrect latency for alcohol (paired t-test, t[23]=0.8482, *p*=0.4051) or sweet milk (paired t-test, t[23]=1.547, *p*=0.1356). HP and LP mice had similar changes in incorrect latency for alcohol (student’s t-test, t[46]=0.6622, *p*=0.5111) or sweet milk (student’s t-test, t[45]=1.105, *p*=0.2752).

Lastly, consumption analysis yielded no differences pre-IA, as HC and LC mice had similar incorrect latencies (student’s t-test, t[46]=0.1394, *p*=0.8898) for alcohol and sweet milk (student’s t-test, t[46]=0.6318, *p*=0.5307). After IA, there were no differences in incorrect latency for alcohol (student’s t-test, t[46]=1.554, *p*=0.1271) or sweet milk (student’s t-test, t[45]=1.077, *p*=0.2874) between HC and LC mice. HC incorrect latency did not change after IA when responding for alcohol (paired t-test, t[23]=0.6598, *p*=0.5159) or sweet milk (paired t-test, t[22]=0.9191, *p*=0.3680). LC mice also saw a strong trend decrease in incorrect latency for alcohol (paired t-test, t[23]=1.889, *p*=0.0716), but not for sweet milk (paired t-test, t[23]=0.5340, *p*=0.5985). HC and LC mice had similar changes in incorrect latency for alcohol (student’s t-test, t[46]=0.9709, *p*=0.3367) or sweet milk (student’s t-test, t[45]=0.0970, *p*=0.9231).

Correct and Incorrect latency results are summarized in **Table 1**. Examining correct and incorrect latency by engagement, preference, and consumption, supports that mice that engage in the 5-Choice display clear stimulus motivational differences that alcohol preference and consumption analyses may miss. Additionally, this data suggests that excessive alcohol intake has a much greater effect in LE mice when responding for alcohol. This may allude to potential development of maladaptive engagement toward alcohol, but further studies are needed for thorough results.

### 3.7 Perseveration in high- and low-engaged mice significantly increases after intermittent access

Excessive alcohol use can lead to maladaptive fixation on alcohol and develop compulsory behavior toward its acquisition (CITE). Perseveration, the continued response toward a stimulus even when it is absent, can be measured by the 5-Choice to elucidate any compulsory-like behavior developing in the mice. Here, we calculated perseverative responses as a percentage as low-engaged mice give lower number of overall responses. Additionally, any mice who did not average 10 correct responses in the last five Late-Stage sessions were excluded as their low response rates would not represent the actual behavior.

Perseveration in HE mice was significantly greater than LE mice when responding for alcohol (**Figure 5A**, Mann-Whitney, *U*=106.5, *p*=0.0487) but not for sweet milk (**Figure 5C**, Mann-Whitney, *U*=168.5, *p*=0.9120). However, after IA, HE and LE mice were similar in perseverative responding for alcohol (**Figure 5A**, Mann-Whitney, *U*=124, *p*=0.1534), as well as for sweet milk (**Figure 5C**, Mann-Whitney, *U*=168, *p*=0.9061). Specifically, LE mice significantly increased perseveration after IA compared to pre-IA (**Figure 5A**, paired t-test, t[14]=0.5.615, *p*<0.0001) where HE mice did not (**Figure 5A**, paired t-test, t[22]=1.537, *p*=0.1387). LE mice had a significantly greater change in perseveration after IA compared to HE mice (**Figure 5B**, student’s t-test, t[36]=2.317, *p*=0.0263). No significant changes pre-IA to post-IA were found for LE (**Figure 5C**, paired t-test, t[14]=1.470, *p*=0.1636) or HE mice (**Figure 5C**, paired t-test, t[22]=0.5312, *p*=0.6006) when responding for sweet milk. There was no difference in the magnitude of change between HE and LE mice for sweet milk (**Figure 5D**, student’s t-test, t[36]=0.6496, *p*=0.5201).

**Figure 5.**
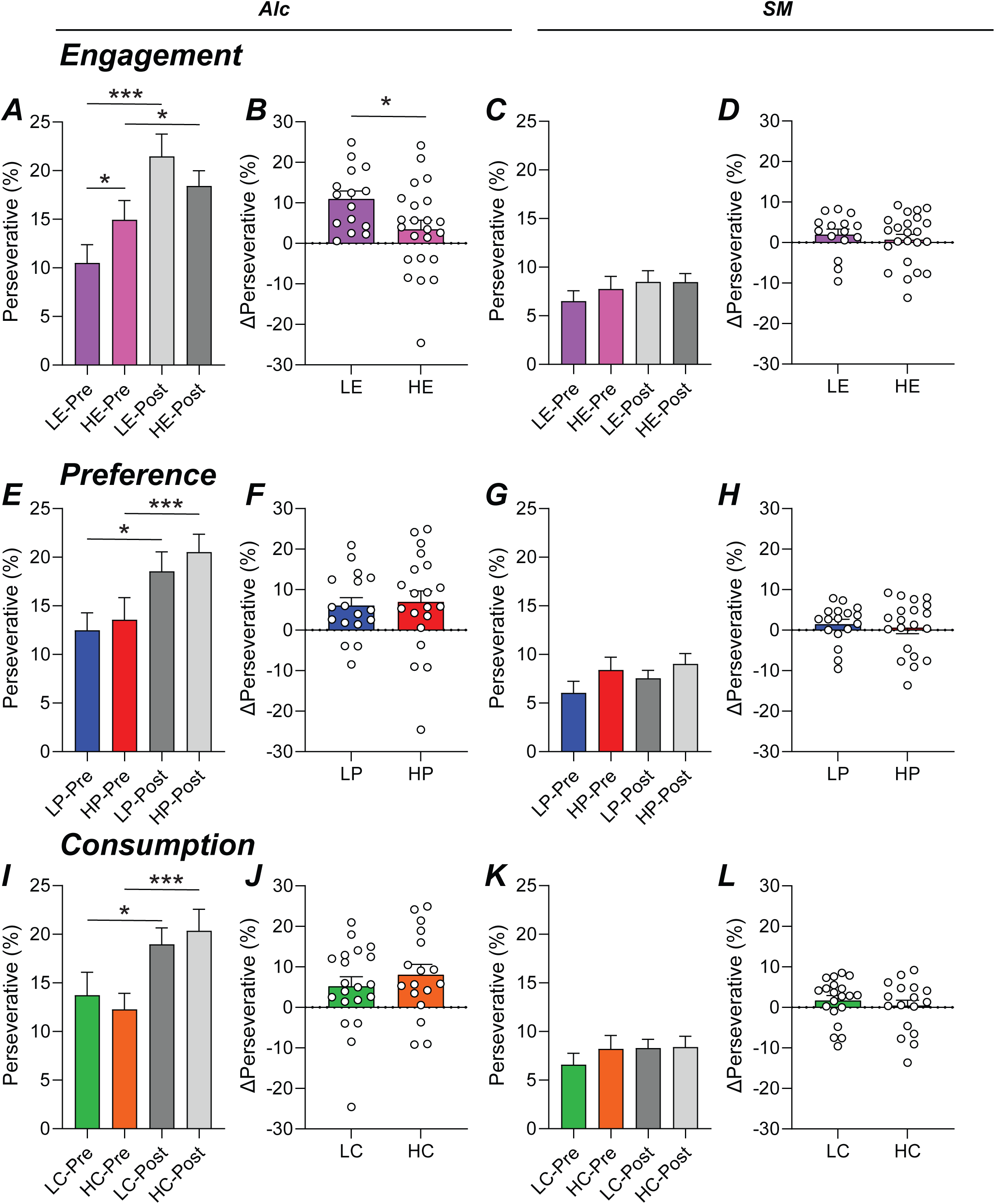
Intermittent access increases perseverative responding for alcohol in low-engaged mice. The percentage of perseverative responding prior to intermittent access was greater in HE mice and became similar between the groups after intermittent access (A). LE mice faced a significantly greater change in perseveration compared to HE mice (B). No changes were seen in sweet milk pre- or post-intermittent access (C) and the overall change remained similar (D). Perseverative percentage analyzed by high and low preference showed increases in perseveration after intermittent access in both groups (E) but no difference in overall change in perseveration between the groups (F). No changes were seen in sweet milk pre- or post-intermittent access (G) and the overall change remained similar (H). Perseverative percentage analyzed by high and low consumption showed increases in perseveration after intermittent access in both groups (I) but no difference in overall change in perseveration between the groups (J). No changes were seen in sweet milk pre- or post-intermittent access (K) and the overall change remained similar (L).

Using alcohol preference-based grouping, there were no differences pre-IA between HP and LP mice when responding for alcohol (**Figure 5E**, student’s t-test, t[35]=0.3649, *p*=0.7174) or sweet milk (**Figure 5G**, student’s t-test, t[35]=1.317, *p*=0.1965). Similarly, there were no differences between the groups post-IA for alcohol (**Figure 5E**, student’s t-test, t[35]=0.7262, *p*=0.4725) or sweet milk (**Figure 5G**, student’s t-test, t[35]=1.065, *p*=0.2941). Both LP (**Figure 5E**, paired t-test, t[16]=3.112, *p*=0.0067) and HP (**Figure 5E**, paired t-test, t[19]=2.541, *p*=0.0199) mice significantly increase perseveration after IA for alcohol but not when responding for sweet milk (**Figure 5G**, LP: paired t-test, t[16]=1.260, *p*=0.2257; HP: paired t-test, t[19]=0.4126, *p*=0.6845). HP and LP mice had similar changes in perseveration for alcohol (**Figure 5F**, student’s t-test, t[35]=0.2560, *p*=0.7994) or sweet milk (**Figure 5H**, student’s t-test, t[35]=0.4487, *p*=0.6564).

Using alcohol consumption classification, there were no differences pre-IA between HC and LC mice when responding for alcohol (**Figure 5I**, Mann-Whitney, *U*=166.5, *p*=0.9222) or sweet milk (**Figure 5K**, student’s t-test, t[35]=0.8974, *p*=0.3757). Similarly, there were no differences between the groups post-IA for alcohol (**Figure 5I**, student’s t-test, t[35]=0.5095, *p*=0.6136) or sweet milk (**Figure 5K**, student’s t-test, t[35]=0.0784, *p*=0.9379). Both LC (**Figure 5I**, paired t-test, t[19]=2.246, *p*=0.0368) and HC (**Figure 5I**, paired t-test, t[16]=3.190, *p*=0.0057) mice significantly increase perseveration after IA for alcohol but not when responding for sweet milk (**Figure 5K**, LC: paired t-test, t[19]=1.424, *p*=0.1708; HC: paired t-test, t[16]=0.1270, *p*=0.9005). HC and LC mice had similar changes in perseveration for alcohol (**Figure 5J**, student’s t-test, t[35]=0.8304, *p*=0.4119) or sweet milk (**Figure 5L**, student’s t-test, t[35]=0.7813, *p*=0.4399).

When comparing perseveration based on engagement, we uncover differences between HE and LE mice prior to IA, unlike analysis based on preference or consumption. Further, the overall change in perseveration of LE mice is significantly greater than HE mice, also not seen in preference or consumption analyses. The behavior of the mice responding for alcohol becomes clearer and may emphasize how LE are more vulnerable to excessive alcohol intake than HE mice.

## 4 Discussion

Alcohol use disorder (AUD) development is heavily promoted by frequent periods of excessive alcohol intake. Individuals with AUD will become abnormally attentive and have high motivation toward alcohol (CITE). Together, attention, motivation, and excessive intake create a strong barrier against treatment and thus it is important to identify weaknesses in this barrier for developing effective therapeutics. The current study is a re-assessment of data previously published that described how alcohol preference was a more sensitive indicator of 5-Choice performance for alcohol compared to alcohol intake (CITE). Additionally, we found that mice were able to adequately learn to the complex 5-Choice task using water restriction and a 10% alcohol reward, a novel method. Interestingly, low-preference mice escalated their performance greatly after intermittent access. These findings correlated 5-Choice behavior to alcohol preference during a 2-bottle Drinking in the Dark paradigm. However, upon second look, we were interested in the innate behavior of each mouse to work for an alcohol reward in the 5-Choice and how excessive intake may enhance this behavior. Importantly, the 5-Choice was used due to its ability to describe various forms of behavioral engagement (e.g., attention, impulsivity, attention) within the same session (CITE). In our current analysis, mice were classified by “engagement” which is the number of correct responses during their final alcohol sessions. Correct responding was chosen to be the best representative example of engagement as it requires the animal to be attentive and allows assessment of motivation as only correct answers deliver the reward and collection latency can be measured. A median split of engagement clearly describes two separate groups. One group, the High-Engaged (HE) display effective learning of the task as accuracy and number of correct escalate as training persists while omission percentage declines. The Low-Engaged (LE) display low, but stable performance throughout all of training, which may suggest that they are not motivated for alcohol enough to work for it. Interestingly, the average correct responses across the Early-Stage sessions correlates with high alcohol preference and consumption, suggesting that preference and consumption will predict greater engagement in the 5-Choice for alcohol. However, across Late-Stage session HE and alcohol preference only hold a strong trend (*p=*0.0500). In Early- and Late-Stage, LE did not correlate to any drinking measure. Similar to our previous finding, alcohol preference and this case, also consumption may be a good predictor of early innate performance for alcohol in the 5-Choice. In relation to humans, this may reflect ones’ drive to put forth effort toward acquiring alcohol, compared to those who will only consume when it is freely available, similar to the LE mice not performing in the 5-Choice but drink during homecage sessions. However, we must further explore the possibility that periodic alcohol sessions promote overall HE effort in the 5-Choice. We would accomplish this by removing Drinking in the Dark sessions so that their only alcohol exposure is within the task.

Sweet milk, the traditional reward (CITE), was used to ensure that mice understood the task. During these sessions mice were no longer water restricted, but rather weight restricted. Our previous work describes the limitations of this (CITE). Briefly, we believe that weight restriction is not appropriate to help motivate for an alcohol reward as alcohol may not be seen as a source of calories for mice. In contrast, we thought weight restriction was appropriate as sweet milk is a great source of calories and satisfies hunger. In future work, only alcohol will be used in the task and will forego sweet milk testing to avoid potential errors. In this work, we felt it necessary to prove that the mice understood the task and were adequately trained. In **Figure 3C**, HE mice showed greater skill in the 5-Choice responding for sweet milk. However, LE mice greatly increased accuracy and correct responding while greatly decreasing omission percentage. Within five sessions, LE mice reached standard criterion for traditional 5-Choice behavior (CITE). We hypothesize that additional sweet milk sessions would show both HE and LE at similar performance. As the HE mice were performing for alcohol the previous 24 sessions (Early- and Late-Stage), we also hypothesize that they have had more time to become more efficient at the task in comparison to LE mice.

The current experiment used the simplest form of the 5-Choice, with a 10s stimulus duration and 5s intertrial interval throughout all of the sessions. Typically, the mice would be trained to reach a stimulus duration between 0.5 and 2s when responding for a sugar-based reward (CITE). Since alcohol is being used, it was unknown how performance would change across a session. “Front-loading” is an aspect of excessive alcohol intake that is described as an escalation of intake within the first 30 minutes of at least a two-hour period (CITE). In our behavioral paradigm, mice are given 30 minutes to respond for alcohol and their behavior may be influenced by the “front-loading” drive. Thus, we desired to make the 5-Choice task simple so we could observe across a single session how alcohol intake may affect performance. Interestingly, **Figure 4E** shows that HE mice decrease motivation for the reward during the second half of their responding. In our previous analysis, we also found this in High-Preference (HP) mice along with having majority of their correct responding occurring in the first half of responding (CITE). This data may reflect a form of cognitive satiety where the individual has the urge to drink but relax after acquiring a sufficient amount of alcohol to eliminate this urge (CITE). Alternately, there is also potential concern that the mice are facing mild cognitive or motor changes. Thus, this study strongly supports a trial-by-trial analysis when using an intoxicant for the reward.

Perseveration, continued screen touches after a correct or incorrect response, is a sign of compulsive-like behavior (CITE). In Figure 5, we analyzed percentage of perseverative responses in all three classifications (engagement, preference, and consumption). Percentage of perseverative responses was used since LE mice had limited overall responding. Additionally, only animals that averaged at least 10 correct responses across their last three alcohol sessions were used in the analysis to reduce low behavior artifacts. Interestingly, only engagement analysis described a difference between HE and LE prior to intermittent access. All analyses observed significant escalation of perseverative of both HE and LE after intermittent access. However, engagement analysis revealed an overall greater change in perseverative responding in LE compared with HE. Sweet milk behavior yielded no differences or changes in behavior between groups or after intermittent access. By focusing on engagement classification, we see that every mouse increased their perseveration after intermittent access of LE mice supporting the danger of excessive alcohol intake in low trait risk individuals. Perseverative responding has been seen to correlate to several brain areas. The orbitofrontal cortex lesions have promoted impulsive and perseverative responding in the 5-Choice during baseline. However, during impulsivity and stimulus challenge testing, premature responding declined while perseveration remained suggesting the orbitofrontal cortex plays a differential role on responding pending difficulty of the task (CITE Chudasama 2003).

Our previous publication describes several limitations of this study (CITE). Briefly, the study does not utilize both sexes. Due to the variability to alcohol preference, consumption, and motivation in mice, we opted to use a large cohort of male mice. Further, having such a large cohort (48 mice) creates an immense amount of work as each mouse needs to be single-housed and trained in the 5-Choice. For example, eight behavioral chambers with 48 mice takes roughly four to five hours every day. In humans, drinking problems in women have risen greatly and may have more inhibitory control issues (CITE Erol and Karpyak 2015, White 2015, Becker and koob 2016, Carvalho 2019, Peltier 2019, Weafer 2015, Nederkoorn 2009). Thus, we understand the importance of potential sex differences and now that our current study has proved mice can be trained with alcohol, every future study should include males and females. Another limitation is that we were unable to obtain blood alcohol levels of the mice after 5-Choice behavior or after homecage drinking. We have previously seen binge drinking levels in C57BL/6 mice reach 1.6g/kg after a 30 min homecage session using 20% alcohol (CITE Lei 2016, Wegner 2017, Kwok 2021), this is roughly 85-100 mg% blood alcohol level. For prospective blood alcohol level after 5-Choice behavior, we can estimate that the mice, on average, had access to 1.3-1.5g/kg of 10% alcohol that, if consumed, may reach a blood alcohol level of 70-80 mg%. As our current experimental design was focused on ability to train mice with alcohol and compare after an excessive intake paradigm, we were unable to plan appropriate times to take blood alcohol levels without potentially effecting ongoing behavior. Though, with the results from this study, we can confidently move forward in collecting blood alcohol levels at specific timepoints.

The purpose of the current study was to emphasize two main concepts. The first concept is how identifying mice by behavioral engagement for alcohol in the 5-Choice delineates at least two different behavior phenotypes. Here, we found that HE early training correlated with alcohol preference and consumption, with preference holding a strong trend during late training. This would suggest that, if screening animals for those that will perform for alcohol in the 5-Choice, you could use homecage drinking as selection criteria to increase the likelihood of greater performing mice. However, as our current design used average preference and intake across training, consistent alcohol homecage drinking may promote 5-Choice behavior. The second concept emphasizes how excessive alcohol intake changes alcohol seeking behavior. Mice with low engagement in the 5-Choice for alcohol throughout Early- and Late-Stage training quickly escalated performance after three weeks of intermittent access, similar to our previous paper where low alcohol preferring mice escalated in their performance. However, by grouping the mice by 5-Choice engagement behavior, the data reflects how intermittent access increases low responding animals in the context of working for alcohol, opposed to correlation of preference and behavior. Thus, we provide a novel analysis method for behavioral engagement for alcohol in the 5-Choice that will prove critical for identifying correlations between behavior patterns and molecular and brain circuits. Altogether, it is important to identify changes in attention, motivation, impulsivity, and compulsivity that may work in concert to drive maladaptive alcohol engagement so that future treatments can be more effective.

## AUTHOR CONTRIBUTION

PS designed the behavioral schedule and collected the data. PS and AS analyzed the data. PS, AS and FH wrote the manuscript. All authors provided critical review of the content and approved of the final version for publication.

## ACKNOWLCDGEMENTS

We would like to thank Hunter Mead and Danielle Maulucci for help with the animal behavior.

---

**Table 1. Response latencies for alcohol and sweet milk by engagement, preference, and consumption classification.**

