## Supplemental Figures for "Engagement for Alcohol Escalates in the 5-Choice Serial Reaction Time Task After Intermittent Access"

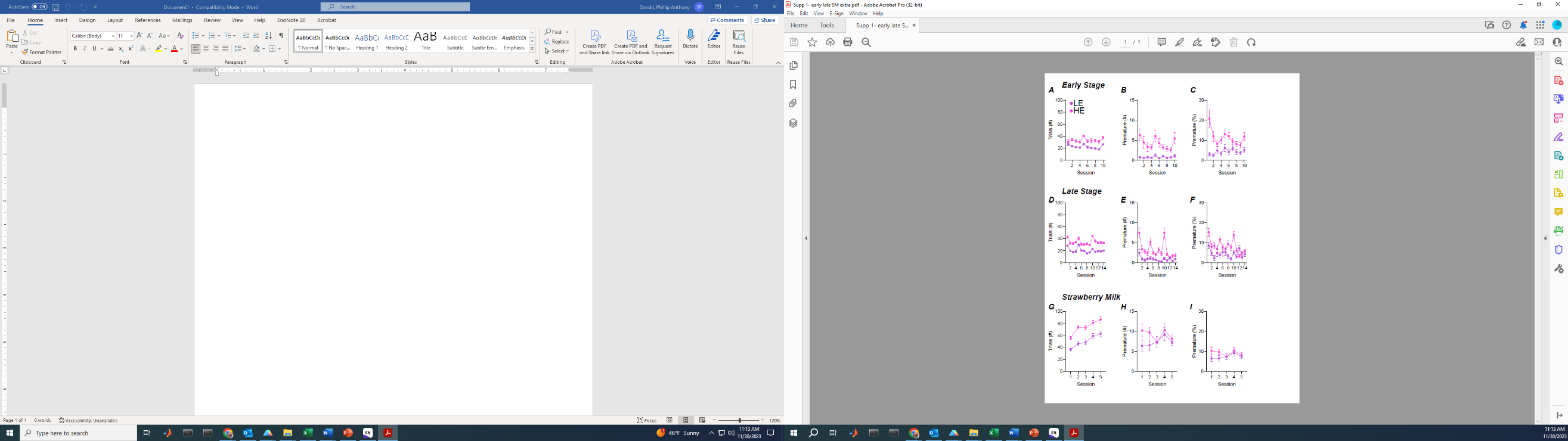

\#

\#

\#

\#

\#

\#

**Supplemental Figure 1. Early, late, and Sweet Milk training in the 5-Choice.** HE mice performed significantly more Trials (A), premature responses (B) and percentage of premature (C) during early stage. HE mice also had greater Trials (D) and premature responses (E) during late stage training, but both had similar percentage of premature responses (F). HE mice performed significantly more Trials (G) during Sweet Milk training, and both groups had similar premature (H) and premature percentage (I).

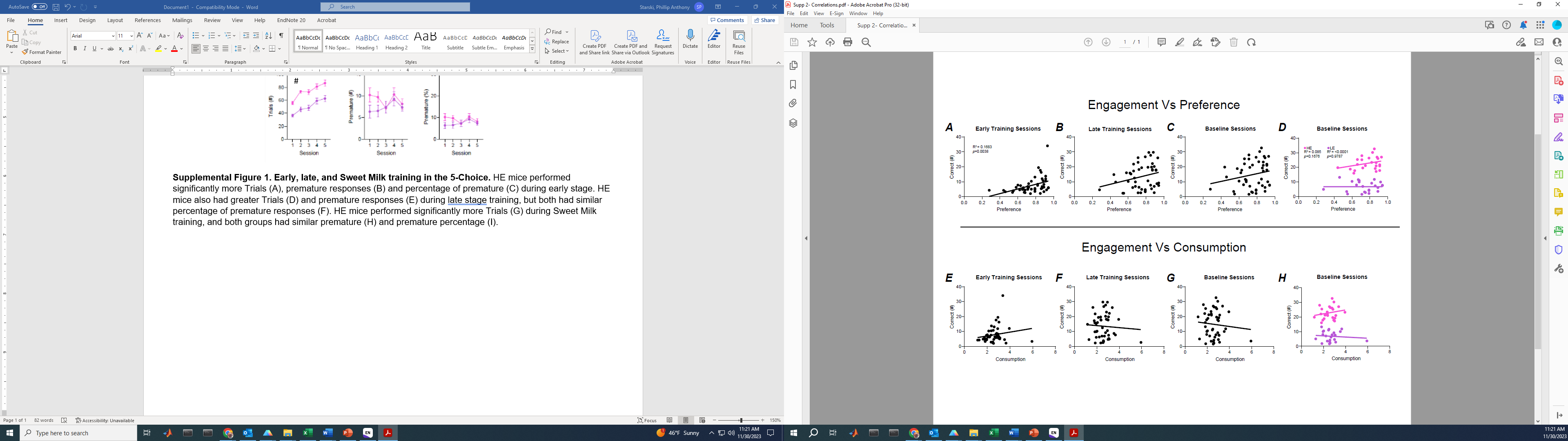

**Supplemental Figure 2. Homecage drinking correlation to engagement performance.** Alcohol preference of all mice correlated with correct responding during early training sessions (A) but dissipated during Late (B) and baseline (C,) sessions, the final five sessions of late stage training. HE and LE mice held no correlation between correct responding and alcohol preference (D). No correlations were found in all mice between correct responding and alcohol consumption for early (E), late (F), or baseline (G) sessions. HE and LE mice held no correlation between correct responding and alcohol consumption (H).
